# Prior-Likelihood Metamers to Distinguish Strategies Underlying Human Bayesian Behaviour in Perception

**DOI:** 10.64898/2026.08.14.744750

**Authors:** Chin-Hsuan Sophie Lin, Nikhita Terence, Marta I. Garrido

## Abstract

Bayesian decision theory proposes that people make statistically rational decisions by combining prior knowledge with sensory information (likelihoods). This framework successfully explains many aspects of human behaviour. However, debate persists over whether people perform precise Bayesian computations (i.e., *explicit Bayesian strategy*) or rely on less demanding strategies—such as approximations or heuristics—that produce Bayesian-like behaviour (i.e., *implicit Bayesian strategy*). To address this, we examined people’s sensitivity to metamers: different prior–likelihood combinations yielding identical optimal policies. An explicit Bayesian observer would show a temporary performance drop immediately after a switch of prior-likelihood combination, followed by recovery, reflecting prior updating. In two studies, we trained participants to estimate hidden target locations drawn from a Gaussian prior. On each trial, scattered dots provided likelihood information. Over time, participants learned the prior and combined it with likelihood information to infer target locations. We then covertly introduced an untrained prior–likelihood metamer. Unlike explicit Bayesian observers, participants’ performance declined after the switch and persisted throughout the untrained pair presentation. This finding challenges strict Bayesian interpretations of task performance and suggests that participants rely instead on *likelihood-sensitive* strategy that is neither explicit Bayesian nor does it not fully integrate prior information. Our study demonstrates how metamer manipulations can distinguish behaviour that merely appears Bayesian, from behaviour genuinely produced by Bayesian computations, and calls for the use of metamers for ruling out alternative explanations of Bayesian-like behaviours.

## Introduction

Humans need to encode sensory information such that the internal representations are coherent with the external world in a manner that is adaptable for survival. Bayesian decision theory (BDT) provides a framework for integrating prior knowledge with new sensory evidence (Berger, 2013; Yuille & Bülthoff, 1996). Rooting in the Helmholtz tradition (Dayan et al., 1995), which stipulates that perception is not a passive copy of the external world but an active, “unconscious inference”, a decision is seen as interpreting uncertain and noisy sensory information to guide behaviour (Berger, 2013; Yuille & Bülthoff, 1996). For each sensory observation, a Bayesian observer integrates a prior over the underlying world state with a likelihood function, the probability of the observation given a hypothesised world state, to infer the posterior probability of possible world states. BDT further denotes that an action is then selected based on a cost (or reward) function defined over possible decision outcomes, allowing the observer to minimise expected cost or equivalently, maximise expected reward, when responding to the inferred world state. Thus, BDT can compute an ideal Bayesian behaviour that serves as a benchmark for statistically optimal decision making. Mounting evidence shows that human behaviour often approximates this optimality by flexibly weighing priors and sensory information (Berniker et al., 2010; Knill & Pouget, 2004; Körding C Wolpert, 2004; Sato & Körding, 2014; Vilares et al., 2012).

The success of BDT has inspired numerous hypotheses about how the brain learns (Bejjanki et al., 2020; Faculty et al., 2020) and represents the world in a Bayesian manner, that is, how it encodes individual elements within Bayesian decision frameworks (Knill & Pouget, 2004; Ma et al., 2006; Pouget et al., 2013; Beck et al., 2008; Ma et al., 2006; Sanborn, 2017; Sanborn & Chater, 2016). Extensive theoretical and empirical work has also sought to explain aberrant behaviours observed in various conditions—from neurodevelopmental disorders (Randeniya et al., 2021), neuropsychiatric conditions (Huys et al., 2015; Valton et al., 2019), ageing (Lin & Faisal, 2018), neuropharmacological manipulations (Vilares & Kording, 2017), to differences between amateurs and experts (Gredin et al., 2023) — proposing that these variations reflect altered Bayesian representations or changed inference processes in the brain. Computational mechanisms behind health conditions have been generated and corresponding treatments proposed by these findings (Huys et al., 2016; Moutoussis et al., 2017). However, despite the considerable advances of the Bayesian brain framework and significant impacts on neuroscientific research, a central question remains: what is the cognitive strategy that supports behaviour appearing Bayesian? Bayes optimal behaviour does not, on its own, imply that the brain uses *explicit Bayesian strategy*: that is, the brain represents each component of BDT and carry out exact inference over them. (Gardner, 2019; Lin & Garrido, 2022; Marr, 2010; Rahnev & Denison, 2018). Indeed, many have argued that given time pressure and cognitive limitations in real-world settings, individuals need to rely on simpler computational strategies to mimic Bayesian optimal behaviour with no representation of individual BDT components (Gigerenzer & Gaissmaier, 2011; Nakayama & Shimojo, 1992; Zhou et al., 2020), which we refer to as *implicit Bayesian strategy*. Sohn and Jazayeri (2021) capitalised on the fact that multiple priors, likelihoods, and cost functions can result in the same optimal Bayesian decision policy and developed an approach for determining if humans derive decisions on separately acquired priors and cost functions in the BDT. The key notion was to check whether participants showed evidence of relearning when unbeknownst to them, one learned prior–cost function pairing (i.e. original) was replaced with a different pair that prescribed the identical optimal decision (i.e. metamer). They tested this idea in two perceptual tasks. In each task, a discrete switch from the original to a novel metamer pair occurred after behavioural performance reached optimality. Using this procedure, they were able to validate an explicit Bayesian strategy in the interval timing task but not in the visuomotor rotation task, demonstrating the effectiveness of metamer designs in disentangling the underlying strategies behind Bayesian behaviours. Given the ubiquity of perceptual decisions and the significance of BDT in neuroscience, it is imperative to assess and understand the underlying cognitive strategies that underpin decision-making. Specifically, various experimental tasks that has been instrumental in formalising Bayesian theories in psychology and psychiatry should be properly scrutinised.

Here, we proposed to expand this “Bayesian metamer” approach, by deriving a “prior-likelihood” metamer to validate whether people perform a Bayesian perceptual task: the coin task (Vilares et al., 2012) using explicit Bayesian strategy. The coin task and its variations (Bejjanki et al., 2016a; Kiryakova et al., 2020a; Sato & Körding, 2014) are among the most commonly used tasks in demonstrating Bayesian behaviour in visual perception. It has been repeatedly shown that during the task people adjust their decision based on the reliability of prior and likelihood information by giving more relative weight to the one with higher precision, a defining characteristic of Bayesian inference. Clinically, the coin task has served as a window to examine how representations of prior and sensory information are affected by neurological (Vilares & Kording, 2017) and psychiatric conditions (Goodwin et al., 2023a; Randeniya et al., 2021). However, the exact cognitive strategy employed for the task, as with many other Bayesian behavioural paradigms, remains unclear. Drawing on previous research, we hypothesised three possible outcomes following a switch to a metamer pair. If participants use an explicit Bayesian strategy, switching to a metamer pair would trigger the learning of a new prior and likelihood, ultimately leading to the identification of the optimal sensory weight. This weight should be the same as that of the original pair (Figure 1.D, dark blue line). A transient decline in Bayesian optimality whilst the new prior and likelihood are learned, followed by a recovery, would be expected (Figure 1.F, dark blue line). If participants employ an implicit Bayesian strategy, the sensory weight (Figure 1D, orange line) and the level of Bayesian optimality (Figure 1.F, orange line) would remain unchanged throughout since the posterior distribution and optimal decision policy yielded by the metamer are identical to those of the original stimulus pair. Finally, if participants employ a likelihood-sensitive strategy, they will focus on learning only the new likelihood as a mean to maintain task performance. Right after the switch, their behaviours should be similar to that predicted by an Explicit Bayesian strategy, with deviations from the original sensory weight and from Bayesian optimality. Overtime, however, limiting learning to the new likelihood leads to a potentially persistent bias in sensory weighting (Figure 1.D, green line) and a prolonged decline in Bayesian optimality (Figure 1.F, green line). To understand the effect of feedback on performance, we compared two experimental designs, with Experiment 1 providing exact location feedback throughout while Experiment 2 providing only partial feedback after prior learning was completed.

**Figure 1.**
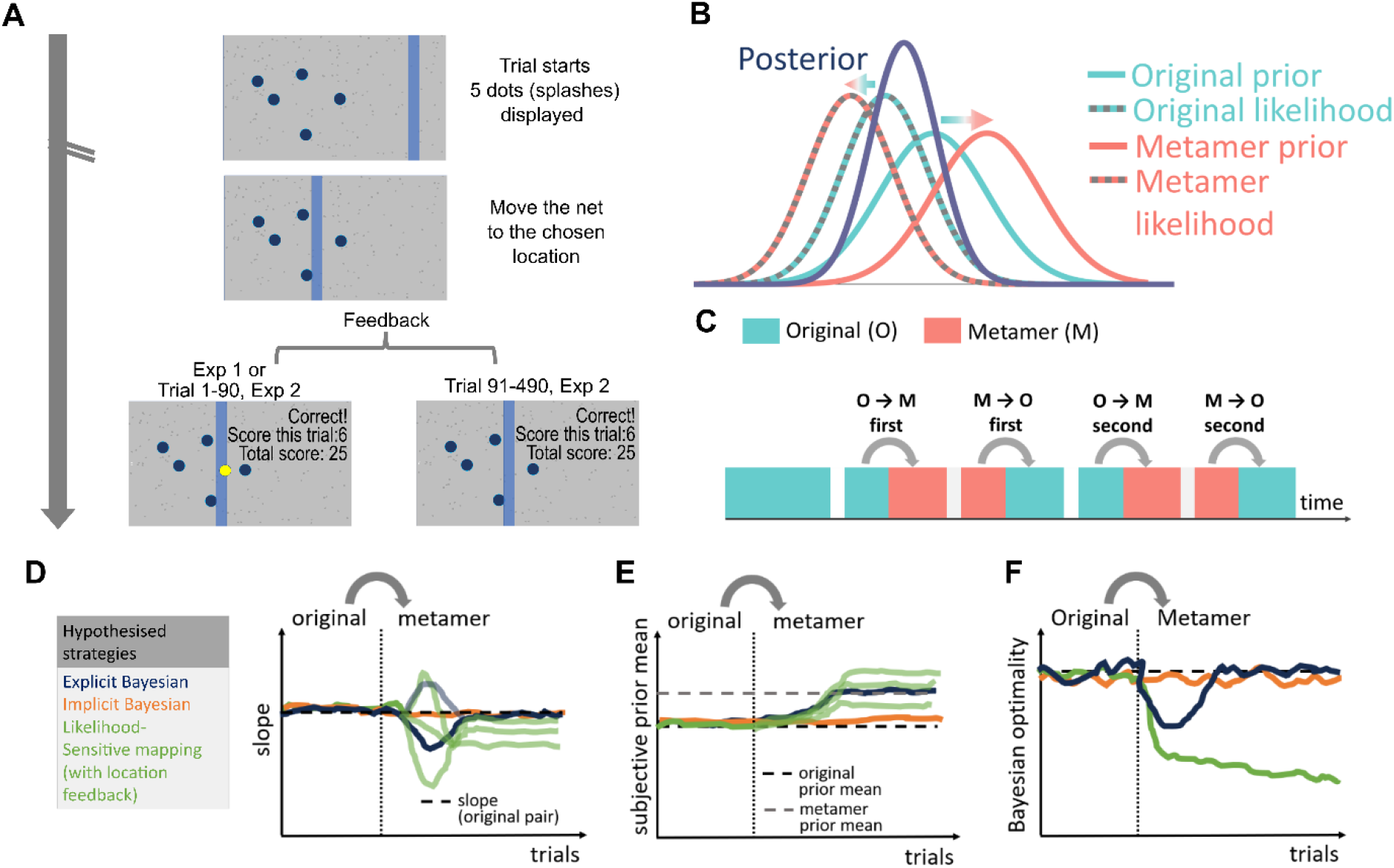
Prior-likelihood metamer in the coin task to test cognitive strategy behind Bayesian behaviours. (A) coin task. Each trial started with 5 dots which represented splashes caused by a thrower throwing a “coin” into the pond (grey screen). Participants were asked to move a blue “net” to the location where they believed the coin has landed (middle panel). Once a response was made, feedback would be displayed for 1-3 second before the next trial started. For experiment 1 and the first 90 trials in the Experiment 2, the true coin location (yellow dot), together with trial score and accumulating score were displayed. For the remaining trials in Experiment 2, trial score and accumulating score were revealed while the yellow dot was no longer presented. (B) Prior–likelihood metamers were pairs of different prior and likelihood distributions that yielded identical posterior distributions and, consequently, the same optimal policy for the coin task. In this study, both priors and likelihoods were Gaussian distributions. The original prior had a mean of 0 and standard deviation (SD) of 0.53; the metamer prior was designed with a mean of 0.036 and SD of 0.53. Because the posterior distribution was known, once the magnitude of the prior mean shift was specified, the corresponding likelihood mean for the metamer pair could be computed (see Methods for detail). (C) Experiment design. Participants first completed the task with the original prior–likelihood pair (green) and thereby learnt the original prior. Subsequently, the task alternated uncued between the original (green) and the metamer (pink) pairs, producing two original→metamer switches and two metamer→original switches in total. (D) Expected sensory weight (slope) under the three hypothesised cognitive strategies. Under an explicit Bayesian strategy, the slope would deviate from the sensory weight for the original pair while the metamer is learned, then recover to the original level once learning is complete (dark blue line). Under an implicit Bayesian strategy, the posterior — and therefore the optimal decision policy — remains unchanged after the switch, so no learning is required and the slope would be indistinguishable between original and metamer conditions throughout (orange line). Under a likelihood-sensitive mapping strategy, participants learn the metamer likelihood; because multiple likelihood–posterior mappings can produce similar reward, the resulting slopes may vary (green lines). (E) Expected subjective prior mean under the three strategies. Under an explicit Bayesian strategy, the subjective prior mean would shift after the switch and later converge on the metamer’s prior (dark blue line). Under an implicit Bayesian strategy, the subjective prior mean should remain unchanged between the original and metamer conditions (orange line). Under a likelihood-sensitive strategy, participants may acquire different subjective priors that yield equivalent reward (green lines). (F) Expected Bayesian optimality (the probability that the observed data match those of an ideal Bayesian observer) under the three strategies. Under an explicit Bayesian strategy, the optimality level is expected to decrease immediately after the switch as participants explore to learn the metamer prior, then rebound after learning (dark blue line). For an implicit Bayesian strategy, the optimality level should remain indistinguishable between original and metamer conditions (orange line). For a likelihood-sensitive mapping strategy, the optimality level would drop from the pre-switch level and fail to recover for a prolonged period (green line).

## Methods

### Participants

The online study was approved by the University of Melbourne research ethics committee (research ethics project reference number 20592) and performed in accordance with the Declaration of Helsinki. Participants were recruited using the University of Melbourne psychology research participation pool. All participants provided written informed consent before data collection. They completed a self-reported questionnaire via Qualtrics online survey platform (www.qualtrics.com) to confirm that they had normal or corrected-to-normal visual acuity, and no history of neurological, psychiatric disorders or substance use. Participants were compensated with course credits for their time. All questionnaire and behavioural data were collected anonymously to protect data privacy.

For Experiment 1, 79 participants (65 female, age mean ± SD = 19.7 ± 2.6 yrs) participated in the experiment. For Experiment 2, 99 participants (71 female, age mean ± SD = 20.0 ± 3.9 yrs) took part. Among them, 4 participants were excluded for having a history of neurological or psychiatric disorders, leaving 95 participants (68 female, age mean ± SD = 19.7 ± 2.7 yrs) for further analyses.

Our sample size was guided both by the first metamer study (which used N = 11) and by a formal power analysis. Using the R package ‘SSDbain’ and the approach described in [42], we estimated that to achieve 80% power to detect an effect in a two-sided t-test at α = 0.08, a minimum of 48 participants would be required.

### Stimuli and apparatus

The experiments were built using the programming software PsychoPy 3 version 2023 and conducted online through the Pavlovia platform. The minimum requirement for screen resolution was 1172 × 553 pixels. It was expected that all participants completed the experiment on a full-screen desktop or personal computer and sat at a comfortable distance away from the screen. All stimuli were set against a grey background and scaled as per the width unit defined in PsychoPy 3. Both experiments took about 1 h to complete.

### Coin Task (Figure 1A)

Detailed descriptions of the coin task can be found in previous publications (Lin et al., 2024a; Vilares et al., 2012). In short, participants were instructed to treat the screen as the surface of a pond. On each trial, a coin (1% screen width in diameter) was thrown into the pond, and participants moved a vertical blue bar (“net”, width 2% screen width) to their estimated coin position. Because the height of the “net” and screen were equal, only the horizontal position was relevant. To induce a prior expectation, participants were informed that the coin thrower initially aimed at a specific target location (the prior mean) for a certain period. After this period, the thrower switched to a new target location. Participants were also told that the thrower might change target locations more than once, thereby implying multiple shifts between the metameric and original prior means. They were also told that the thrower did not have perfect aim but the level of accuracy (how closely the coin landed to the intended target) remained consistent throughout (the prior standard deviation (SD) *σ_P_* was 5.3%, which was not revealed to participants outright but had to be learned). While the coin location was not visible at the time of the estimation, its “splashes” were visible through five blue dots (1% screen width in diameter), to provide the sensory information (or likelihood) for the coin’s location. The horizontal locations of these five dots were drawn from a Gaussian distribution with a mean of the horizontal coin position in that trial and an assigned standard deviation of 0.94% screen width. The likelihood SD *σ_L_ wa*s thus 0.94/√5.

For each trial, participants would score between zero and ten points. Ten points were awarded if the centre of the coin exactly overlapped with the centre of the net. Scores decreased linearly as the distance between the centre of the coin increased and became zero when the net no longer overlapped with the coin. Actual coin locations and scores were displayed throughout Experiment 1. In Experiment 2, coin locations were shown during the initial 90 trials while participants learned the distribution of the original pair (Figure 1A) to ensure that Experiment 2 participants’ prior learning experience matched that of Experiment 1.

Thereafter, coin location feedback was removed to minimise the chance of reinforcement-based learning. However, feedback on the points scored was still provided at the end of each trial to maintain participant engagement.

### Computing prior-likelihood metamers

The original prior was a Gaussian distribution with a mean *μ_Po_* of 0 (the horizontal extent of the screen ranged from −0.5 to 0.5, with 0 denoting the midline of the screen) and a SD *σ_Po_* of 0.053. We held the SDs of the prior *σ_P_* and likelihood *σ_L_* distributions constant throughout (i.e. *σ_P_*_o_ = *σ_P_*_m_ and *σ_Lo_* = *σ_L_*_m_). We shifted the prior mean by 0.67 SDs, so the mean (μ_Pm_) of the metamer prior was 0.036 (Figure 1B). Given this change in prior mean, for the posterior to remain identical to the original, the corresponding metamer likelihood mean *μ_Lm_* was determined as follows: Once the level of prior mean change has been decided, the corresponding of the metamer likelihood mean could be computed using the following equation:

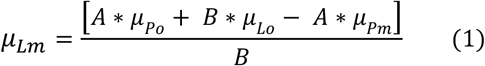

where

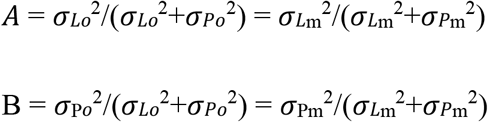

### Experimental procedure details (Figure 1C)

Participants completed 450 trials in Experiment 1 and 490 trials in Experiment 2. In both experiments, the first 90 trials comprised the original prior–likelihood pair which aimed to induce prior-learning. Participants took roughly 30 seconds to 1-minute breaks every 90 trials to avoid fatigue and maintain engagement. Four alternating uncued transitions between the original and metamer pairs took place at trial 131, 221, 311, 401, yielding two original→metamer (trial 131 and 311) and two metamer→original (221, 401) switches (Fig. 1.C). We chose switch trials such that they were not immediately before or after breaks, in order to avoid any confounding effects introduced by rest periods. The task was self-paced and participants completed the experiment within 60 minutes.

Feedback differed between the two experiments. In Experiment 1, participants received both the coin location and the trial score on every trial. In Experiment 2, the coin location was shown only for the first 90 prior-learning trials (Fig 1.A); all subsequent trials provided only the trial score. As mentioned, the change aimed to minimise error-based learning following each prior–likelihood pair switch.

### Data analysis

All analyses were conducted in RStudio (R version 4.4.2), with the significance level set at *p* < .05. Data normality was evaluated using the Shapiro–Wilk test. Performance differences between conditions were assessed using a repeated-measures ANOVA when data were normally distributed. When the normality assumption was violated, a linear mixed model was first used. Post hoc pairwise comparisons were performed with Tukey’s HSD adjustment.

Where appropriate, Bayes factors were reported to quantify evidence in favour of the null hypothesis. We quantified between the original-metamer pairs and across switches how closely participants’ performance matched that of Bayesian optimal observers using three measures— slope, subjective prior mean, and optimality index. We also examined whether instantaneous and average scores, a reward-based behavioural marker, changed between the original-metamer pairs and across switches. Below we described the rationale behind choosing these measures and how we computed each in detail.

#### Estimating likelihood reliance (slope) and subjective prior mean using BDT

In the coin task, integrating prior and likelihood information according to the reliability of each is considered the hallmark of ideal Bayesian behaviour. Analytically this is evaluated by fitting a regression line predicting the estimated coin location (i.e., the chosen net location) from the centre of the splashes (i.e. the likelihood) (Vilares et al., 2012).

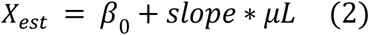

The slope of the regression line quantifies how much participants relied on the likelihood. A slope between 0 and 1 indicates that both likelihood and prior information influenced net placement: values closer to 1 reflect greater reliance on the likelihood, while values closer to 0 indicate greater reliance on the prior. Based on the Bayes’ rule, an optimal net location *Χest* is

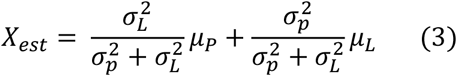

This means that for a Baye-optimal observer, the slope is expected to equal 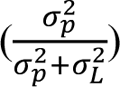 and to be 0.61 for both original and metamer pairs in the current study. We first inspected whether slopes from the original pair matched predictions of the ideal-observer model. Slopes during metamer trials were then compared against those of original trials in attempts to differentiate the hypothesised strategies participants likely employed: If participants were explicit Bayesian learners, we expected a transient deviation from the values of those observed in the original pair immediately after a switch, followed by a rapid return (Figure 1D blue line). By contrast, implicit Bayesian learners (Figure 1D orange line) were expected to maintain slopes indistinguishable from the original pair. For likelihood-sensitive observers, there may be multiple linear mappings between the likelihood and the selected locations that produce equally satisfactory participant scores; consequently, the slopes may remain the same or differ across mappings (Figure 1D green lines).

To characterise the temporal evolution of slope values, for each trial, we regressed chosen net locations on splash centres using a sliding window of the following twelve trials. When fewer than twelve trials remained in one pair, we progressively reduced the window size, stopping once fewer than six trials were available. The final five trials of each trial type were thus excluded from the figure. We conducted repeated-measures analyses to test whether, after the first couple of trials following each switch, slope values on metamer trials were indistinguishable from those on original trials, as would be predicted for both explicit and implicit Bayesian learners. Since the first 90 trials of each experiment constituted the prior learning phase and our previous work showed that slopes stabilised after approximately 50 trials following a change of prior (Lin et al., 2024a), we excluded the initial 90 trials of each experiment plus the first 50 trials following each switch from this statistical analysis.

Explicit Bayesian observers aim to estimate likelihood and prior distributions accurately; therefore, assessing both the slope and the subjective prior mean provides a fuller measure of how closely participants’ behaviour conformed to the predictions of an explicit Bayesian strategy. Looking at equations (2) and (3), we can see that under the Bayesian scenario, the intercept *β*_0_ in equation (2) equals (1-slope) * *μ_P_*, meaning that we can estimate subjective prior mean using the following equation:

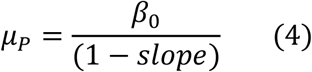

We examined the time course of inferred prior means using a sliding window identical to that employed in the slope analysis. For the statistical analysis, we took trials the same as those included in the slope statistics. It should be noted that slope values approaching 1 produce arbitrarily large prior mean estimates. These undesirable artefacts likely arise from noise introduced by the fitting procedure and/or measurement error. For each condition, participants whose slopes exceeded 0.95 were excluded from the prior mean calculations.

Outliers in the prior mean estimates were then removed from the statistical analyses using a 3 MAD criterion. Like for slope, we expect that with an explicit Bayesian strategy, the subjective prior mean shifts after the switch and gradually converges on the metamer’s prior as it is learnt (Figure 1E, blue line). With an implicit Bayesian strategy, the subjective prior mean would remain unchanged between original and metamer conditions (Figure 1E, orange line). With a likelihood-sensitive strategy, participants would acquire distinct subjective priors that nonetheless yield comparable reward (Figure 1E, green lines).

#### Optimality index (OI)

Adapted from (Acerbi et al., 2014), for every horizontal screen position **χ**, we can calculate the probability of hitting the true coin location *р_hit_* given that a participant places the net at position **χ**. The analytical solution for *р_hit_* based on the mean ***μ*** and standard deviation ***σ*** of the true Gaussian posterior for each prior–likelihood combination, is

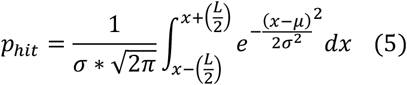

where *L* is the effective width of the capture region, defined as the sum of the net width *l* and the coin diameter *d*. The hitting probability is maximal (i.e. max (*p*_ℎ*it*_)) when **χ** = ***μ***. The optimality index (OI) is defined as *p_int_(x_net_)/max(p_hit_)*.

The OI represents the probability that the behavioural responses were generated by Bayes-optimal observers. It is more sensitive than slope, prior mean or score alone, for tracking the optimality of the decision policy utilised by people, which is exactly the measure needed to differentiate cognitive strategies with metamer paradigms (Sohn & Jazayeri, 2021) and refer to Figure 1.D). Additionally, OI is computed on a trial-by-trial basis, as opposed to slopes and inferred prior mean, which require integration across multiple trials or restriction to a subset of data (such as excluding slope > 0.95 trials when computing subjective prior mean). Thus, as we shall see in the results section, OI provides a more stable measure for evaluating the time course of Bayesian optimality in a dynamic environment, such as our design. To distinguish between explicit Bayesian, implicit Bayesian and likelihood-sensitive strategies, we compared OIs in pre-switch, early post-switch and late post-switch windows for each switch (i.e. −25 to 0, 1 to 25 and 26 to 50 trials, where 0 denotes the time of the switch). This approach follows Sohn and Jazayeri (2021), who showed that explicit Bayesian learners typically exhibit an immediate dip in optimality after the switch followed by recovery after approximately 25 trials. Thus, OIs in the early post-switch window should be lower than both the pre-switch and the late post-switch OIs for explicit Bayesian learners (Figure 1F, blue line). Implicit Bayesian learners, by contrast, would show little or no difference in OIs across the three windows (Figure 1F, orange line). Likelihood-sensitive learners would display a prolonged reduction in optimality, so both early and late post-switch OIs are expected to be lower than pre-switch OIs (Figure 1F, green line).

#### Mean score

For each trial, participants received a score from 0 to 10 based on the horizontal distance between the coin and the net, with higher values indicating better performance. We sought to determine whether participants scored similarly for original and metamer trials once their behaviours had stabilised following an original->metamer switch. Explicit and implicit Bayesian learners would be expected to exhibit mean scores and levels of optimality comparable to original trials in the later metamer trials. (Sohn & Jazayeri, 2021). Examining scores can also reveal incomplete metamer learning: if participants’ scores remain unchanged immediately after the switch, they may adopt a suboptimal but “good-enough” policy. In contrast, lower scores combined with persistently reduced optimality would support the likelihood-sensitive strategy. Trial-wise group-averaged scores were thus calculated to capture both the immediate impact of the switch on participants’ reward-based performance and the time course over which performance evolved. Mean scores of each condition (pair and switch) after discarding the very first 90 trials in each experiment plus the first 50 trials after each switch were computed before statistical analyses.

## Results

### Slope and subjective prior mean

In order to disambiguate amongst plausible rational decision-making models that combine prior knowledge with sensory information (likelihoods) we first asked whether there was a slope (sensory-prior relative weighing) and prior mean change between original and metamers. In Experiment 1, slopes from 71 participants were retained (8 of 79 excluded) after applying the 3-MAD criterion. In Experiment 2, slopes from 73 participants were retained (22 of 95 excluded) according to the same criterion. For the prior-mean analyses, a higher proportion of participants were excluded owing to the two-tier exclusion criteria detailed in the Methods. In Experiment 1, the numbers retained per condition were: original-switch 1, 51; original-switch 2, 50; metamer-switch 1, 48; metamer-switch 2, 37, corresponding to 28, 29, 31 and 42 exclusions, respectively, all out of 79. In Experiment 2, the numbers retained per condition were: original-switch 1, 54; original-switch 2, 60; metamer-switch 1, 49; metamer-switch 2, 51, corresponding to 41, 35, 46 and 44 exclusions, respectively, out of 95.

We first assessed whether responses to the original stimulus pair matched Bayesian predictions prior to the initial switch. The observed median slopes were 0.66 (95% CI [0.57, 0.73]) for Experiment 1 and 0.80 (95% CI [0.70, 0.86]) for Experiment 2. In Experiment 1 the slope did not differ significantly from the optimal slope of 0.61 (bootstrap p = .23), but in Experiment 2 it was significantly higher than the veridical value (bootstrap p < .001).

Overall, participants qualitatively followed the Bayesian pattern of sensory weighting, although they showed an over-reliance on the likelihood in Experiment 2, consistent with numerous previous coin-task studies. (Bejjanki et al., 2016b; Goodwin et al., 2023b; Kiryakova et al., 2020; Lin et al., 2024). Subjective prior means were 0.008 (95% CI [-0.002 0.020], bootstrap p = .16 against actual prior mean) in Experiment 1 and 0.012 (95% CI [0.006 0.019], bootstrap p < .001 against actual prior mean) in Experiment 2. This slight rightward bias relative to the actual prior mean is also consistent with our previous findings (for a meta-analysis, see (Zhao et al., 2025)). Having established that participants exhibited Bayesian-like behaviour, consistent with previous research, we then compared the slopes and subjective prior means between the original stimuli and their metamers to determine the cognitive strategies employed.

Slope values began to decrease approximately 30 trials after the switch from original to metamer pairs. Although they rebounded and stabilised during later metamer trials, they remained lower than those for the original pairs, particularly during the second switch (Figure 2A, top panel). This was supported by a linear mixed model (Figure 2A, lower panel) revealing significantly lower slopes for metamer pairs (median ± IQR = 0.51 ± 0.73) than for original pairs (median ± IQR = 0.65 ± 0.53; main effect of pair, p = .004). Slopes were also lower in the second switch (median ± IQR = 0.57 ± 0.68) than in the first switch (median ± IQR = 0.68 ± 0.49; p = .008; Figure 2A, inset of lower panel). There was no significant interaction between pair type and switch (p = 0.16).

**Figure 2.**
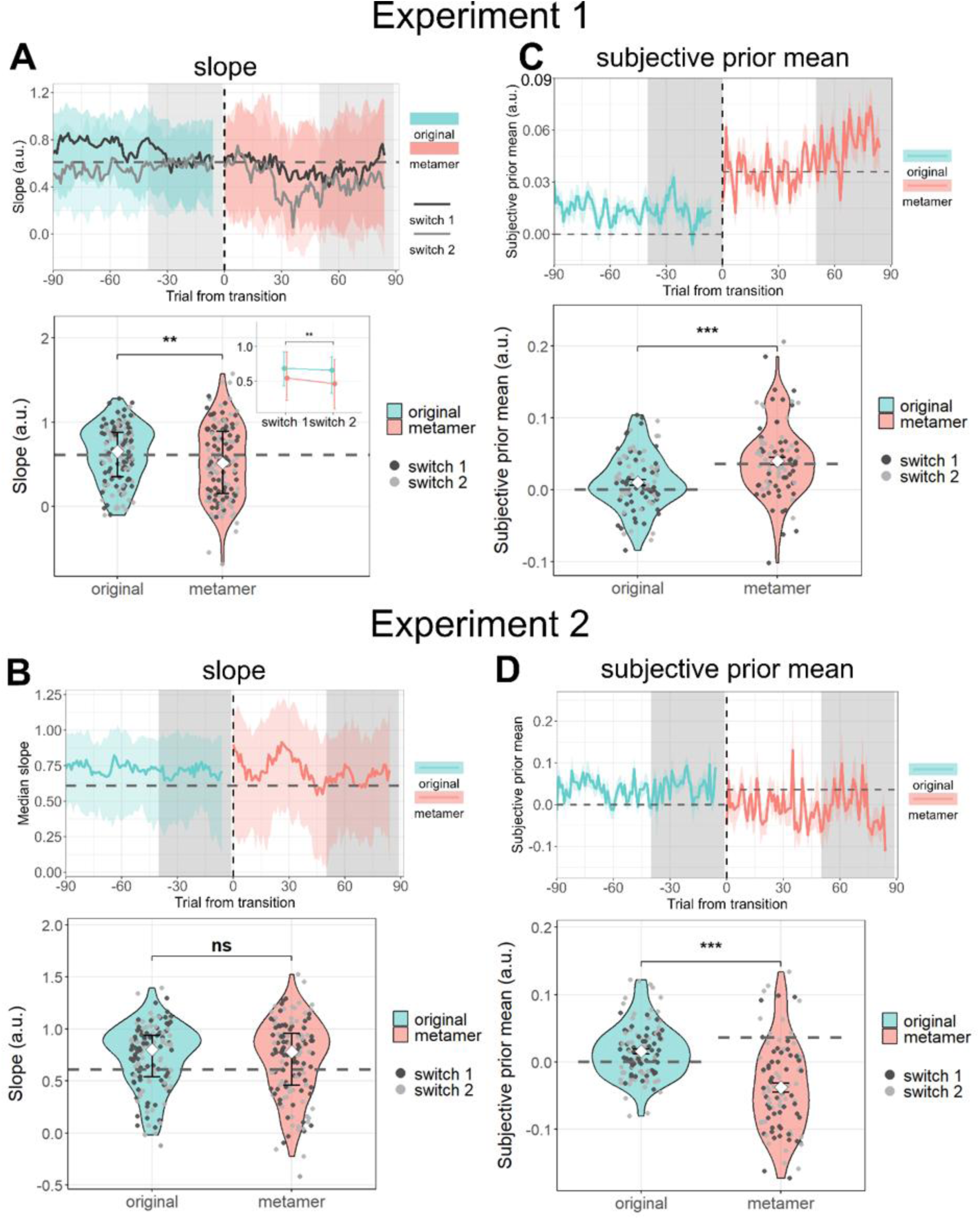
BDT parameters: slopes and subjective prior means (time courses and grouped data presented in violin and scatter plots). (A) Experiment 1. Upper Panel shows the time course of the slope. There was a decrease after each switch to metamer trials, reaching a trough at ≈30 trials post-switch, followed by a partial rebound and stabilisation by ≈50 trials. The degree of change was larger in the second switch. Dash horizontal line indicates the optimal slope. Shaded grey areas indicate the trials used for statistical analyses and shown in the violin plots and white shaded areas indicated trials excluded for those analyses and plots; the same convention applies to all time-course panels. Violin plots show that slopes on metamer trials were significantly lower than on original trials (median ± IQR: metamer = 0.51 ± 0.73; original = 0.c5 ± 0.53; p = .004). The inset shows that slopes in the second phase were also lower than in the first phase (median ± IQR: second = 0.57 ± 0.c8; first = 0.c8 ± 0.4S; p = .008). (B) Upper panel: In Experiment 2, slopes increased shortly after the switch but subsequently returned to values similar to those on the original trials. The dashed horizontal line marks the true prior mean for both the original and the metamer. Violin plots indicate no significant difference in slope between original and metamer trials (linear mixed model, p = .80). (C) Experiment 1. Upper panel: Subjective prior means shifted rightwards immediately following the switch. The dashed horizontal lines mark the true prior means for both the original and the metamer. Violin plots show subjective priors on metamer trials were significantly more right-leaning than on original trials (mean ± SE: metamer = 0.042 ± 0.00c; original = 0.010 ± 0.004; p < .001), consistent with the shift in the true prior. (D) Experiment 2. Subjective prior means on metamer trials shifted leftwards relative to original trials, i.e. in the direction opposite to the true prior shift (mean ± SE: metamer = −0.038 ± 0.007; original = 0.01c ± 0.004; p < .001). Note: Time courses were computed using a sliding window of the following 12 trials. When fewer than 12 trials remained for a given pair, the window size was reduced stepwise; no estimate was computed when fewer than six trials were available. Therefore, the final five trials of each condition were omitted from the plotted data. Significance markers: ** p < .01; *** p < .001

Trial-wise subjective prior mean in the Experiment 1 immediately moved towards the actual metamer prior mean after the switch (Figure 2C upper panel). A repeated-measure ANOVA confirmed this observation: there was a significant pair main effect (F (1,49) = 18.04, p < .001) with the subjective prior mean of metamer moving in line with the direction of actual prior mean shift, albeit numerically larger than the true mean of 0.036 (mean ± s.e. = 0.042 ± 0.006, 95% CI [0.031 0.053]). There was no significant effect of switch (F (1,49) = .26, p = .77) or interaction (F (1,49) = 0.15, p = .45). Taken together, these changes in slope and prior mean following the switch were incompatible with the predictions of an implicit Bayesian strategy. Although the rebound in slope and the update of prior-mean estimates were qualitatively consistent with predictions of explicit Bayesian observers, the significant decrease in slopes, the larger drop during the second switch in particular, means we cannot rule out the possibility of a likelihood-mapping strategy.

In Experiment 2, the instantaneous slope values markedly increased following the switch from original to metamer stimuli but subsequently returned to values comparable to those observed in original trials (Figure 2B. top panel). A linear mixed model revealed no significant main effects or interaction for slopes (pair: p = .59; switch: p = .20; interaction: p = .86, Figure 2B. lower panel), again a picture of possibly either explicit Bayesian or likelihood-sensitive strategies. However, the subjective prior mean, while changing right after the switch (Figure 2D, top panel), moved towards the opposite direction of the true prior mean shift (repeated-measure ANOVA F (1,59) = 37.38, p < .001; mean ± s.e.: metamer = −0.038 ± 0.007 and original = 0.016 ± 0.004), meaning participants failed to learn the metamer prior mean. These findings from both experiments again casted doubts on the explicit Bayesian strategy.

For both Experiments 1 and 2, a substantial number of participants were excluded from the calculation of the subjective prior mean based on the 3MAD outlier criterion. To assess robustness, we re-analysed data without the 3MAD exclusion (excluded by the “slope > 0.95” and “slope < 0” criteria only) and obtained qualitatively identical results to the data presented above. Thus, these results also ruled out an implicit Bayesian strategy and casted doubt on the plausibility of an explicit Bayesian strategy (Figure S1): In Experiment 1, the subjective prior mean of metamer trials was significantly more rightward than on original trials (A linear mixed model, pair main effect p <.001); median ± IQR: metamer = 0.048 ± 0.110, original = 0.009 ± 0.062. In Experiment 2, the prior mean shifted leftward in the metamer (median ± IQR = −0.033 ± 0.114) compared with the original trials (median ± IQR = 0.017 ± 0.062). Although the linear mixed model did not detect a significant difference (pair-type main effect: p = .07), Bayes factor overwhelmingly suggested the alternative hypothesis (BF_10_ = 5.5 × 10^3^).

### Optimality Index (OI)

Since based on the slope and subjective prior mean data we could not disambiguate between explicit Bayesian and likelihood-sensitive strategies we turn to the optimality index (OI) metric which has distinct predictions for both (see Figure 1F).

No OI outliers were excluded based on the 3MAD criterion in either Experiment 1 or 2. In Experiment 1 (Fig 3A), a linear mixed model showed a highly significant main effect of pair type (F(2, 390) = 93.69, *p* < .001): median OI values are lower in the metamer trials (median ± IQR = 0.28 ± 0.27) compared to the original trials (median ± IQR = 0.43 ± 0.31). There was no significant switch main effect (*p* = .89), but a significant interaction (F(2, 390) = 3.79, p = .024). Further examinations showed that the pair effect was significant for all three types of switch trials (pre-switch [−25 to 0], early post-switch [1 to 25], and late post-switch [26 to 50] windows, where 0 corresponds to time of the switch; all p < .001). Thus, the significant interaction arose because the original–metamer difference was largest in the pre-switch period (Δ = −0.15) and was slightly attenuated in the post-switch periods (Δ = −0.08 early; Δ = −0.10 late). Within each pair type, OI did not differ significantly across phases (all Tukey-corrected p > .11), confirming that the interaction reflects a change in the size of the pair-type effect rather than a directional shift.

**Figure 3.**
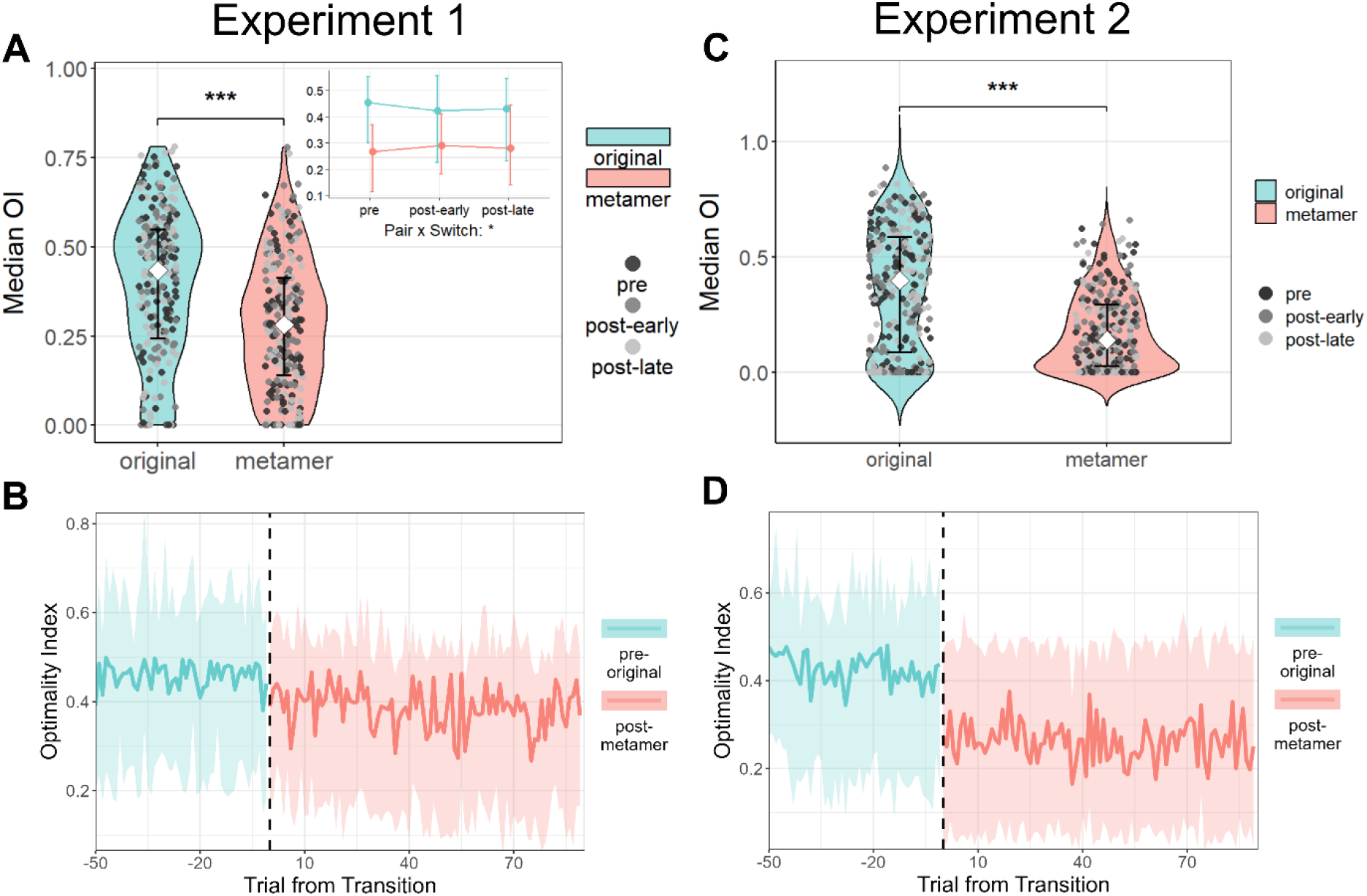
Optiamlity index from Experiment 1 (A. and B.) and Experiment 2 (C. and D.) (A and C) Violin plots show original (green) versus metamer (red) optimality index. Scatter dots represent participant level data (black dots: pre-switch −25 to 0, grey dots the early 25 post-switch and light grey the 2c-50 post-swiitch trials, where 0 correspond to the switch time. For both Experiments, OIs of metamer trials were significantly lower than that of original trials. In Experiment 1, the difference between original and metamer in the pre-switch trials were larger than those in the post-switch trials (3.A inset) (B and D) Panels B and D plot the trial-by-trial optimality index from the 50 pre-switch original trials (green line) through the S0 post-switch metamer trials (red line), averaged across participants and across both switches. In both experiments, the OI of metamer trials after switch (vertical dash line) was consistently lower than those of original pre-switch trials, and there was no evidence of systematic improvement or deterioration in OI over the post-switch period.

To determine whether participants sought to acquire a posterior matching the metamer, as predicted for explicit Bayesian learners, we examined the temporal evolution of post-switch metamer trials. We fitted a mixed-effects model to compare the average OIs from the first ten and the last ten metamer trials. There was no significant difference between the first and last ten trials (F (1, 78) = 0.71, *p* = .40, BF_10_ = 0.16), suggesting that optimality did not systematically increase within a given post-switch period. There was likewise no significant effect of switch number (first versus second switch; F (1, 78) = 0.009, p = .93) nor significant interaction (F (1, 78) = 0.05, p = .82), showing that this absence of within-period improvement in metamer-trial OI was consistent across switches. The trial-wise trajectories of the OI, shown in Figure 3B, mirrored this pattern. These persistently reduced OIs after switch supported a likelihood-sensitive strategy.

The OIs pattern of Experiment 2 was very similar to that of Experiment 1 (Fig 3C & D), but presented with an even more pronounced decrease in OIs of metamer trials. A linear-mixed model showed a significant main effect of pair type (F(1, 470) = 299.33, p < .001), with the original pair yielding higher OIs than the metamer. Neither the main effect of switch (F(2, 470) = 0.40, p = .668) nor the pair × switch interaction (F(2, 470) = 0.90, p = .408) reached significance, indicating that the pair-type difference in optimality was stable across the pre-switch, early post-switch and late post-switch periods. A linear mixed model comparing the first ten versus last ten metamer trial OIs again found no significant difference between the first and last ten trials (F (1, 94) = 0.19, *p* = .66, BF_10_ = 0.16). Whilst there was no significant main effect of number of switches (first versus second switch, *F* (1, 94) = 2.84, *p* = .09), BF₁₀ = 1.72 showed weak evidence favouring the alternative hypothesis: The median OIs of metamer trials decreased slightly from the first switch (0.17 ± 0.38) to the second (0.13 ± 0.33), also in favour of the likelihood-sensitive strategy. There was no interaction (*F* (1, 94) = 1.83, *p* = .18, BF₁₀ = 0.10).

### Mean score

In Experiment 1, mean score among 72 participants (7/79 excluded) were kept for further analyses based on 3MAD criterion. Trial-wise group-averaged scores (Figure 4A) and violin and scatter plots of mean scores (Figure 4B) showed no difference between original and metamer trials. Mean scores were 0.57 ± 0.03 (mean ± s.e.) among original and 0.54 ± 0.03 (mean ± s.e.) among metamer pairs. A repeated-measures ANOVA conducted to evaluate the effects of pair (Original vs Metamer) and switch (1^st^ vs 2^nd^) found no significant main effects or interaction (pair: F (1,71) = 0.44, p = .51; switch: F (1,71) = 1.29, p = .26; interaction: F (1,71) = 0.003, p = .96). A Bayesian analysis found moderate evidence for the absence of a pair (BF_10_ = 0.16) or switch (BF_10_ = 0.22) main effect.

**Figure 4.**
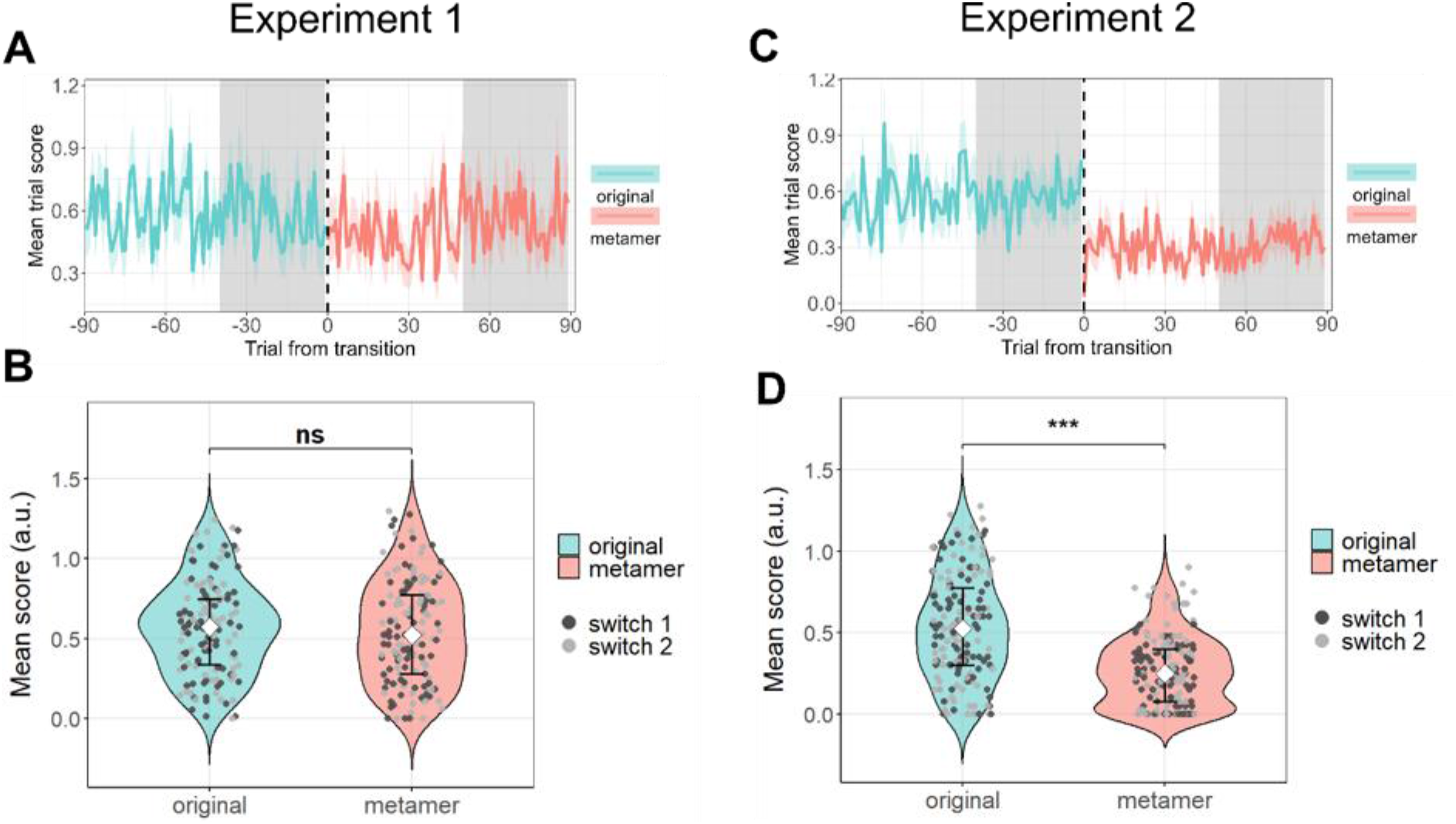
Trial-wise averaged scores and mean scores and trial-wise grouped average scores. (A) Trial-wise grouped average scores for Experiment 1, comparing original (green) and metamer (red) trials. Grey shaded areas indicate the trials included in the statisticl analyses and presented in violin/scatter plots; white areas indicate trials excluded. The same shading convention is used in (C). (B) Mean scores for Experiment 1. There was no significant difference between original and metamer trials (p = 0.51). (C) Trial-wise grouped average scores for Experiment 2, using the same shading convention as in (A). (D) Mean scores for Experiment 2. Mean score on metamer trials was significantly lower than on original trials (p < 0.001; median ± IQR: original = 0.53 ± 0.48; metamer = 0.25 ± 0.33).

In Experiment 2, mean score among 84 participants (11/95 excluded) were kept for further analyses based on 3MAD criterion. Figure 4C showed that scores reduced immediately after the switches to metamer trials and remained persistently lower than those of original trials. A linear mixed model confirmed the pattern and found a significant main effect of pair (F (1,249) = 96.24, p < .001). There was no significant switch main effect (F (1,249) = 1.48, p = .23) or interaction (F (1,249) = 2.21, p = .14; BF_10_ = 0.16). Medians collapsed across switches (Figure 4D) showed participants scored higher during the original trials (median ± IQR = 0.53 ± 0.48) than metamer trials (median ± IQR = 0.25 ± 0.33).

Together, these findings indicate that although participants in Experiment 1 appeared to acquire a decision policy that preserved reward-based performance, that policy depended on trial-by-trial location feedback. Therefore, in the absence of such feedback in Experiment 2, participants were unable to recover performance to pre-switch levels, supporting a likelihood-sensitive strategy.

## Discussion

Bayesian decision theory (BDT) has been highly successful in describing perceptual decision-making behaviours. However, behaviour that appears Bayesian does not necessarily demonstrate that the brain uses explicit Bayesian strategies: i.e. explicitly represents components of the Bayesian equation or performs exact Bayesian computations (Bowers & Davis, 2012a; Jones & Love, 2011). Similar behavioural outcomes can arise from error-based implicit Bayesian strategies (Gardner, 2019; Gigerenzer & Brighton, 2009; Sohn & Jazayeri, 2021) or approximate algorithms (Sanborn, 2017). Studies have attempted to differentiate between these strategies but have been limited by assumptions that Bayesian agents have accurate representation of priors and likelihoods (Beierholm et al., 2009; Kiryakova et al., 2020a). To address this, we used the metamer paradigm (Sohn & Jazayeri, 2021) in the coin task (Vilares et al., 2012) to test whether people form and maintain explicit representations of Bayesian model components (explicit Bayesian strategy), or instead manifest Bayesian-like behaviours that emerge from another strategy.

We first examined slopes and subjective prior means. Although slopes and subjective prior means matching BDT predictions do not prove an explicit Bayesian strategy, we tracked these measures across metamer trials because an explicit Bayesian observer would attempt to re-estimate the prior and likelihood; even if participants cannot fully achieve optimal. In Experiment 1, while we found that the subjective prior mean changed in the direction of the true prior mean shift (Figure 2.C), slope, as a proxy of likelihood precision estimation, deviated from the true probability distribution (Figure 2.A). This pattern is inconsistent with an implicit Bayesian strategy (Figure 1D & E), but it does not clearly distinguish between an explicit Bayesian strategy and a likelihood-sensitive strategy.

The optimality index (OI) allowed us to distinguish among the candidate strategies. In Experiment 1, OIs showed a small but significant reduction immediately after the switch from the original to the metamer pair, and this reduction persisted throughout the post-switch period (Figure 3A and B). Such a decline is inconsistent with an implicit Bayesian account, which predicts little or no change in OI after the switch (Figure 1F, orange line). An explicit Bayesian model predicts an initial dip in OI followed by recovery as observers learn the metamer prior; the absence of any rebound therefore contradicts the explicit Bayesian prediction (Figure 1F, blue line). By contrast, a likelihood-sensitive mapping strategy predicts a sustained reduction in OI due to formation of a new likelihood–policy mapping, and our results are consistent with this hypothesis (Figure 1F, green line).

However, could it be possible that participants, while executing explicit Bayesian strategy, took a ‘optimally lazy’ (Acerbi et al., 2017) approach? That is, they ceased to learn further since rewards (i.e. scores) did not differ. Afterall, the decrease of OIs in Experiment 1, although significant, were only modest. The score also did not decrease at all when the switch to the metamer pair occurred (Figure 4A). In Experiment 2, we removed trial-by-trial coin-location feedback. The primary aim of this manipulation was to minimise error-based learning. However, a natural consequence of this design is that maintaining performance becomes more demanding, meaning that any evidence of prior learning would be all the more pronounced should participants be operating as explicit Bayesian learners. Contrary to the explicit-Bayesian prediction, we observed a marked and sustained drop in mean score.

Participants failed to acquire the information required to sustain performance during metamer trials. Again, we observed a significant and consistent drop in OIs following the switch from original to metamer trials, consistent with the predictions of a likelihood-sensitive mapping strategy. In fact, the subjective prior mean shifted slightly away from, rather than toward, the true prior mean.

People thus seemed, in Experiment 2, to choose a conservative strategy: exploiting the same likelihood-sensitive mapping despite scoring feedback clearly informed a change of environment. The dynamics of optimality index further proved the idea: a quantitatively more significant drop in the optimality index was identified throughout the metamer trials with no improvement at any part during the whole Experiment. Taking findings from both Experiments together, we showed while participants displayed qualitatively Bayesian behaviour for both original and metamer, detailed analysis supported participants relied on forming a likelihood-sensitive strategy, possibly by creating a mapping between likelihood and posterior, rather than forming and maintaining representations of Bayesian model components.

Sohn and Jazayeri (2021) previously designed prior-cost function metamer pairs, demonstrating that metamer can be an effective way of disentangling the cognitive strategies underlying Bayesian behaviours. They suggested that similar designs could be applied to prior-likelihood pairs. To our knowledge, our study is the first empirical investigation to construct a prior-likelihood metamer to validate a Bayesian visuospatial task. We demonstrated the generalisability of the metamer approach, which allowed us to directly test whether observers maintain explicit representations of priors and likelihoods or rely on other learning mechanisms, thereby advancing our understanding of the principles governing human perceptual decision-making. We were able to do so without employing numeric modelling, further showing the power of the metamer paradigm.

Thus, we argue for the value of conducting metamer validation tests when evaluating commonly used Bayesian tasks or before deploying new Bayesian paradigms.

We chose the coin task because the task and its variants (Bejjanki et al., 2016; Sato & Körding, 2014; Vilares et al., 2012; Berniker et al., 2010; Kiryakova et al., 2020) have been widely used in evaluating perceptual inference. This task, by consistently demonstrating that people’s behaviours matched the pattern of integrating prior and likelihood information in a qualitatively Bayesian-like fashion, has been considered a hallmark of Bayesian observers. Studies, including some of our own (Goodwin et al., 2023; Randeniya et al., 2021), have also used this task to investigate altered Bayesian model representations in neurodivergent populations. We have argued for the need not to conflate Bayesian-fashioned behaviours with explicit Bayesian computations (Lin & Garrido, 2022). To this end, we previously applied a transfer measure in the coin task: testing how well people generalised learned prior information to new scenarios to validate computation behind the coin task (Lin et al., 2024). Those findings suggested that people use probabilistic recognition models to generate an approximately optimal posterior representation. However, in our previous work we acknowledged that the transfer design could not clearly distinguish an explicit-Bayesian strategy from a likelihood–sensitive strategy The results of this study indicated that participants likely employ a likelihood-sensitive strategy. The practical implication is clear: the coin task may not be the best option for understanding prior learning and representation in either neurotypical or neurodivergent populations. The finding does not invalidate past discoveries employing the coin task. However, it echoes arguments made by many researchers (Bowers & Davis, 2012b; Griffiths et al., 2012; Rahnev & Denison, 2018) that researchers should be aware of the limitations of Bayesian decision theory and clearly state the difference between computationally and algorithmically Bayesian behaviours (Lin & Garrido, 2022) when describing the goals and outcomes of studies.

It may seem paradoxical that the true posterior stayed the same throughout, despite a clear decline in the OI for the metamers. This is because our study created a shift of likelihood mean as a corresponding change to the prior mean shift that was outside the sampling space during the original phase. As such, a likelihood-posterior approximator did not sufficiently cover the mapping between new likelihood distribution and posterior (Dasgupta et al., 2020). Future studies could attempt to construct metamer pairs such that the original likelihood encompasses the metamer likelihood, so that the metamer likelihood–posterior mapping is already sampled during original trials. Under such a design, any immediate change in optimality following the introduction of metamer trials would reflect an attempt to learn the underlying components of BDT.

Bayesian decision theory has been influential in shaping our understanding of perception and biological computation. However, there is increasing consensus that equating Bayesian behaviours with explicit Bayesian computation in humans is misleading and counterproductive. By employing a prior-likelihood metamer paradigm in an extensively applied perceptual task, we identified the strategy underlying seemingly Bayesian behaviour, demonstrating that metamer design provides a powerful tool for discriminating between strategies that produce Bayesian-like outcomes.

## Supplementary Information

The supplementary information of this manuscript is available online at https://figshare.com/s/05f635a3a244da0bd952

## Acknowledgments

We are grateful to Evie Mallinos for her help with recruiting and running participants.

## Open Practices Statement

Data and code will be available on https://github.com/sophietwlim/Metamer upon the publication of the manuscript. CSL can also be contacted for questions about data and code. None of the experiments was preregistered.

